# Female LSD1 Conditional Knockout Mice Have an Increased Bone Mass

**DOI:** 10.1101/2022.06.25.497607

**Authors:** Kristina Astleford-Hopper, Elizabeth Bradley, Kim C. Mansky

## Abstract

Osteoclasts are large multinucleated cells that degrade bone mineral and extracellular matrix. Investigating the mechanisms by which osteoclast differentiation is epigenetically regulated is a key to understanding the pathogenesis of skeletal related diseases such as periodontitis and osteoporosis. Lysine specific demethylase 1 (LSD1/KDM1A) is a member of the histone demethylase family that regulates gene expression via the removal of mono-and dimethyl groups from H3K4 and H3K9. Prior to our study, little was known about the effect of LSD1 on skeletal development and osteoclast differentiation. Here we show conditional deletion of LSD1 in the myeloid lineage results in a decrease in osteoclast differentiation and activity. Furthermore, Lsd1cKO female, but not male, mice have an increased bone mass. Lastly, we demonstrate that LSD1 can form a complex with CoREST, HDAC1 and HDAC2 suggesting a mechanism by which the combination of methylation and acetylation of histone residues regulates osteoclast gene expression.

## Introduction

Osteoclasts are large multinucleated cells responsible for resorbing bone mineral and extracellular matrix by secretion of catalytic enzymes and acids such as tartrate resistant acid phosphatase and cathepsin K [1]. Osteoclasts play an important role in the bone remodeling process by breaking down worn out, or broken bone and subsequently sending signals to osteoblasts to lay down new bone mineral [2, 3]. This delicate balance of bone resorption coupled with bone formation during remodeling is essential to maintain skeletal homeostasis. An imbalance in this process ultimately leads to bone disorders such as osteoporosis and Paget’s disease [4]. Currently, RANKL inhibitors (e.g., Denosumab) and bisphosphonates are the primary drug treatments used to treat osteoporosis; however, they associate with adverse effects, specifically microfractures, atypical femoral fractures and osteonecrosis of the jaw [5, 6]. The development of new drugs specifically targeted may ultimately lead to better treatment of osteoporotic patients. To create these new treatments, it is important to better understand how genes within osteoclasts are regulated. Epigenetic regulators represent major orchestrators of gene expression control; thus, elucidation of their functions within osteoclasts may yield new therapeutic avenues [7].

Epigenetics is the study of changes in gene expression that can be heritable or a result of environmental factors that do not change DNA sequences themselves [8]. The main forms of epigenetic modifications are histone protein modifications, DNA methylation, and short non-coding RNA sequences [7]. The histone proteins H2A, H2B, H3 and H4 can all be modified via addition or removal of acetyl, methyl, phospho, sumo or ubiqityl groups to either upregulate or downregulate gene expression [8]. One major benefit to studying epigenetic regulators is that they are reversible modifications thus making them desirable targets for drug therapies [8]. In fact, many drugs targeting epigenetic regulators have already been developed, FDA approved and shown to be clinically effective [8]. Thus, the importance of studying these epigenetic regulators is quite relevant.

Lysine specific demethylase 1 (LSD1 or KDM1A) is a member of the histone demethylases that functions by removing mono-or di-methyl groups from histone 3 lysine 4 (H3K4) or histone 3 lysine 9 (H3K9) [9]. LSD1 as well as LSD2 differ from the other members of the demethylase family as they require FAD binding to perform their demethylase activity.

Removal of methyl groups from H3K4 results in a decrease in gene expression whereas removal of methyl groups from H3K9 results in an upregulation of genes, exemplifying that LSD1 has two very distinct roles in regulating gene expression [9]. LSD1 has already been well studied in many different cancers such as breast and prostate cancer; therefore, drugs that target LSD1 have already been developed and are being tested in human subjects [10-13]. However, it is currently unknown whether these drugs would be beneficial in other disease models such as osteoporosis. Prior studies established that conditional deletion of LSD1 in osteoblasts results in increased osteoblast differentiation and mineralization ability via the upregulation of WNT7B and BMP2 expression [14]; however, the role of LSD1 in osteoclast differentiation and activity is currently unknown.

In this study, we sought to understand how a conditional deletion of LSD1 effects osteoclast differentiation. We show that deletion of LSD1 using the LysM-cre, targeting cells in the myeloid lineage, results in a decrease in osteoclast differentiation and activity, and results in an increase in bone mass in mice. These results, together with previously published work in osteoblasts, suggest that LSD1 inhibitors may be a beneficial treatment for patients that suffer from osteoporosis.

## Materials and Methods

### Ethics

All animal experiments were performed in accordance with institutional guidelines by the Institutional Animal Care (IACUC, protocol# 2104-39006A) and the Committee of the Office for the Protection of Research Subjects at the University of Minnesota, Minneapolis.

### *Lsd1* mice

Mice were housed in groups of 4-5 with access to standard rodent chow and water at all times. The housing room is maintained on a 14:10 hour light:dark cycle with controlled temperature and humidity. *Lsd1* floxed mice in a C57BL/6 background were obtained from Jackson Laboratories (Bar Harbor, ME) via Dr. Stuart Orkin’s laboratory (Harvard University, Boston, ME). *Lsd1* floxed mice were crossed with B.6129-*Lyzstm1(cre)lfo/J* mice (LysM-cre) (Jackson Laboratories) which have Cre-recombinase expressed in myeloid cells, which includes osteoclasts. Genotyping primers for Lsd1^fl/fl^ mice listed in **Table 1**.

**Table 1:**
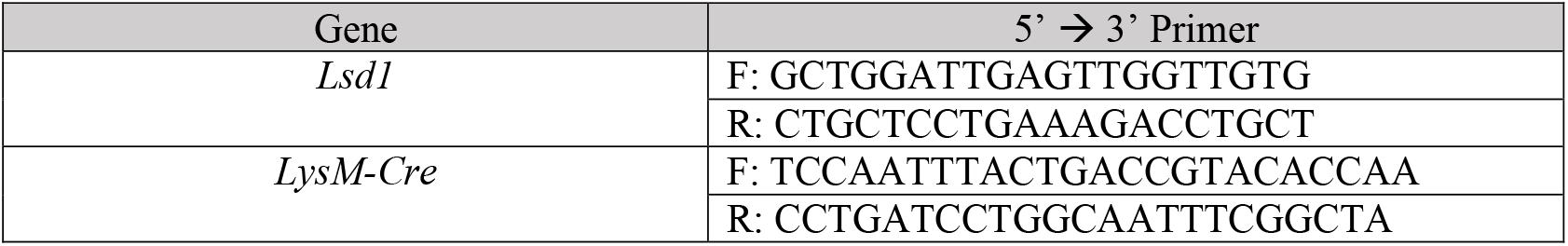
LSD1^fl/fl^ mouse genotyping primers

### In vitro osteoclast analyses

#### Primary osteoclast culture

Femora and tibiae were dissected from Lsd1^fl/fl^ LysM-cre-(Lsd1WT) and Lsd1^fl/fl^ LysM-cre+ (Lsd1cKO) and all adherent tissues were removed. Femora were saved for *in vivo* analyses. Primary bone marrow monocytes from isolated from the tibiae by flushing the marrow cavity with osteoclast media (phenol red-free alpha-MEM (Gibco) with 25 units/mL penicillin/streptomycin (Invitrogen), 400 mM L-Glutamine (Invitrogen), and 5% fetal bone serum (Atlanta Biologicals). Red blood cells from the marrow sample were lysed using a Red Blood Cell lysis buffer (10 mM KHCO_3_, 150 mM NH_4_Cl, 0.1 mM EDTA pH 7.4). The remaining cells were cultured in 10 cm tissue culture dishes overnight (TPP, MidSci) in osteoclast media supplemented with 1.5% CMG 14-12 (cell culture supernatant containing M-CSF). Non-adherent cell populations in the supernatant were collected and replated in 12-well cell culture plates (TPP, MidSci) at a concentration of 2×10^5^ cells per well in osteoclast media supplemented with 1.5% CMG. Cell cultures were subsequently supplemented every two days with 1.5% CMG and 5 ng/mL of RANKL (R&D Systems) to induce osteoclastogenesis.

#### Tartrate-resistant acid phosphatase (TRAP) and DAPI staining

Differentiated osteoclasts were rinsed with 1X DPBS (Gibco) and fixed with 4% Paraformaldehyde (Thermo Scientific) for 20 minutes. Osteoclasts expressing TRAP were stained using the Naphthol AS-MX phosphate and Fast Violet LB salt protocol (BD Biosciences Technical Bulletin #445). Osteoclasts were imaged and photographed using light microscopy. Images were analyzed using NIH ImageJ to measure the number and size of TRAP positive osteoclasts. Cells were then stained with 0.1% DAPI dye in PBS for 10 minutes in the dark at room temperature. Cells were imaged and photographed using fluorescent microscopy. Nuclei were quantified using NIH ImageJ.

#### Bone resorption assay

Primary bone marrow monocytes were plated on Osteo Assay surface plates (Corning) at a concentration of 100,000 cells per well and allowed to fully differentiate. Cells were initially plated with osteoclast media supplemented with 1.5% CMG and subsequently given osteoclast media at a pH of 6.8 supplemented with 1.5% CMG and 5 ng/mL of RANKL every two days.

On day 7 of osteoclast differentiation after RANKL stimulation, media was aspirated off and 10% bleach was added to each well and allowed to sit for 10 minutes at room temperature. The bleach solution was removed, and each well was washed with dH_2_O twice and allowed to dry at room temperature. Plates were imaged and photographed using light microscopy. Resorption pits were analyzed using NIH ImageJ.

#### Real time quantitative PCR analysis

RNA was isolated and purified using RNA Plus Mini Kit (Qiagen) and quantified using a nanodrop spectroscopy. cDNA was prepared from 1 ug of purified RNA using the iScript cDNA Synthesis Kit (Bio-Rad) as stated in the manufacturer’s protocol. Each reaction contained 500 nM of both the forward and reverse primers, 10 μl of iTaq Universal Sybr Green Supermix (BioRad), 8.8 μl DEPC H_2_O (Ambion) and 1 μl of cDNA. The RT-qPCR protocol is as follows: 95°C for 3 minutes, 40 cycles of 94°C for 15 seconds, 58°C for 30 seconds, and 72°C for 30 seconds. Melting curve analysis followed: 95°C for 5 seconds, 65°C for 5 seconds, and lastly 65°C to 95°C with 0.5°C increments for 5 seconds each. The forward and reverse primer pairs for each gene are shown in **Table 2**.

**Table 2:**
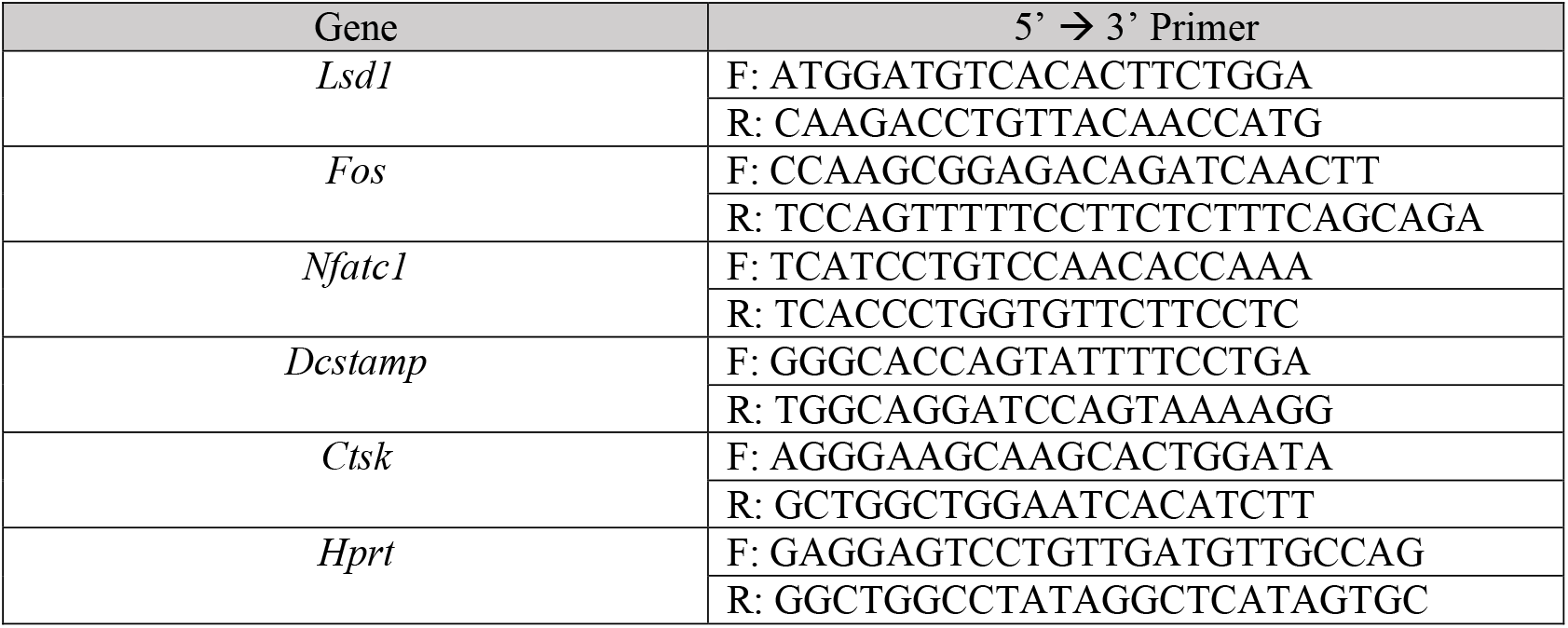
RT-qPCR primers

#### Immunoblotting

Protein lysates were harvested from osteoclasts using modified RIPA buffer (150mM NaCl, 0.25% sodium deoxycholate, 50 mM Tris pH 7.4, 1% IGEPAL, 1mM EDTA) supplemented with Halt Protease & Phosphatase Inhibitor Cocktail (Thermo Scientific). Lysates were purified by centrifugation at 13,000 rpm at 4°C. Proteins were resolved by SDS-PAGE and transferred to PVDF membrane (Millipore). Membranes were incubated overnight with primary antibodies (Antibody list shown in **Table 3**). HPR-conjugated anti-rabbit or anti-mouse (GE Healthcare) was incubated with membranes, washed, and developed using Western Bright Quantum (Advansta) detection agent. Images were acquired using BioRad Chemitouch.

**Table 3:** Immunoblotting antibodies

| Antibody | Source | Catalogue Number |
| --- | --- | --- |
| Anti-LSD1 | Abcam | Ab17721 |
| Anti-Mono-methyl Histone 3<br>Lysine 4 | Abclonal | A2355 |
| Anti-Mono-methyl Histone 3<br>Lysine 9 | Cell Signaling | 14186S |
| Anti-HDAC1 | Cell Signaling | 5356S |
| Anti-HDAC2 | Cell Signaling | 5113S |
| Anti-Acetylated Histone 3<br>Lysine 9 | Cell Signaling | 9649S |
| Anti-Alpha-Tubulin | Cell Signaling | 2144S |

#### Immunoprecipitation

Bone marrow monocytes were isolated as described above. Non-adherent cell populations in the supernatant were collected and replated in 10 cm cell culture plates (TPP, MidSci) at a concentration of 5×10^6^ cells per plate in osteoclast media supplemented with 1.5% CMG. Cell cultures were subsequently supplemented every two days with 1.5% CMG only or 1.5% CMG and 30 ng/mL of RANKL to induce osteoclastogenesis. Cells were harvested by scaping cells off plates with PBS. Cells were then lysed using Pierce IP Lysis Buffer (Thermo Scientific) supplemented with Halt Protease & Phosphatase Inhibitor Cocktail. Lysates were purified by centrifugation at 13,000 rpm for 10 minutes at 4°C. 10% of the lysate was collected for IP input and the rest of the lysate was split up evenly to be incubated with 5 μg of antibody overnight.

EZview Red Protein G beads (Sigma-Aldrich) were washed and centrifuged at 5,000 rcf for 1 minute twice. Lysate and antibody mixes were then allowed to rotate and incubate with beads for 2 hours at 4°C. Beads were centrifuged at 5,000 rcf for 1 minute and supernatant was removed. Beads were subsequently washed with PBS and centrifuged thrice. Immunoblotting using the beads diluted in PBS was performed following the above protocol.

### In vivo bone analyses

#### Micro computed tomography analysis

3-month-old femora were isolated, wrapped in gauze, and stored in PBS at -80°C. At the time of scanning, femora were defrosted to room temperature and scanned in PBS with a 1 mm aluminum filter using a XT H 225 micro-computed tomography machine (Nikon Metrology Inc., Brighton, MI, USA) at an isotropic voxel size of 7.11 μm. Scan settings were set to 120kV, 61μA, 720 projections, 2 frames per projections and integration time of 708 milliseconds. 3D reconstruction volumes were made for each scan using CT Pro 3D (Nikon Metrology Inc., Brighton, MI, USA). These 3D volumes were then converted to bitmap datasets using VGStudio MAX 3.2 (Volume Graphics GmbH, Heidelberg, Germany). Scans were rotated using DataViewer (SkyScan, Bruker microCT), prior to analysis. SkyScan CT-Analyzer was used to perform Morphometric analyses following Bruker’s instructions and guidelines for the field (PMID: 20533309). The trabecular bone analysis was performed in the distal metaphysis starting 0.5 mm proximal to the growth plate and extended 1.5 mm proximally toward the diaphysis. The cortical bone analysis was a 0.5 mm section at the mid-diaphysis. Automated contouring was used in the region of interest for both trabecular and cortical bone with some manual editing as needed for each sample. Global thresholding was used to remove surrounding tissue from analysis in both 3D trabecular and 2D coritcal analyses.

#### Paraffin-embedded section staining

3-month-old femora were isolated and fixed in Z-fix (Anatech LTD) and placed in 10% EDTA (pH 7.4) for decalcification, paraffin-embedded sectioning, and histological staining. Bone sections were then de-paraffinized using xylenes and rehydrated via an ethanol gradient. Sections were then TRAP stained using the above protocol at 37°C for 1 hour. Sections were counterstained with methyl-green for 15 seconds and prepared for imaging using permount mounting media, cover slipped and allowed to dry for 24 hours. Images were taken using light microscopy and analyzed using NIH image J.

#### Statistical analysis

All results are expressed as means with standard deviations. For all *in vitro* experiments, graphs represent an average of at least three independent experiments performed in triplicate. The *in vivo* data represents all the samples gathered graphed together. None of the samples were removed as outliers. Student t-tests were used for all experiments. All statistical analyses were performed using GraphPad Prism 8.

## Results

### Female Lsd1cKO mice have increased bone mass

The role of LSD1 in regulating osteoblast differentiation and activity has been studied; however, its role in osteoclasts is unknown. To study the role of LSD1 in regulating osteoclast differentiation, we created a mouse model by crossing *LysM-cre* mice with *Lsd*^*fl/fl*^ mice to create *Lsd1*^*fl/fl*^*;LysM-cre*^*-*^ (herein referred to as Lsd1WT) and *Lsd*^*fl/fl*^ *LysM-cre*^*+*^ (herein referred to as Lsd1cKO) mice. 3-month-old mice were used to determine the *in vivo* skeletal phenotype of male and female Lsd1cKO mice. 3D reconstruction and volumetric analysis using micro-computed tomography (μCT) revealed that Lsd1cKO female, but not male mice, have significantly more trabecular bone volume to total volume (BV/TV, 0.8955±0.2367 percent increase) compared Lsd1WT mice (**Figure 1A-B, G-H**). Furthermore, Lsd1cKO females but not males have significantly more trabeculae (Tb.N, 0.2492±0.05295 increase), less trabecular spacing (Tb.Sp, 0.01924±0.004754 decrease), and a higher connective density (Conn.Dn, 51.48±11.53 increase) (**Figure 1C-F, I-L**) compared to Lsd1WT mice. When analyzing the cortical phenotype of these mice, the female Lsd1cKO mice have a significant increase in mean total cross sectional tissue area (T.Ar, 0.06841±0.03216 increase) and tissue perimeter (T.Pm, 0.1347±0.06188 increase). (**Figure S1E & F**), however neither the female nor male mice have any changes in cortical BV/TV suggesting that they have no significant change in their cortical phenotype (**Figure S1A-D, G-L**). Together, these results suggest that Lsd1cKO female mice have increased bone mass, but the loss of Lsd1in the myeloid lineage does not significantly affect the male skeleton.

**Figure 1.**
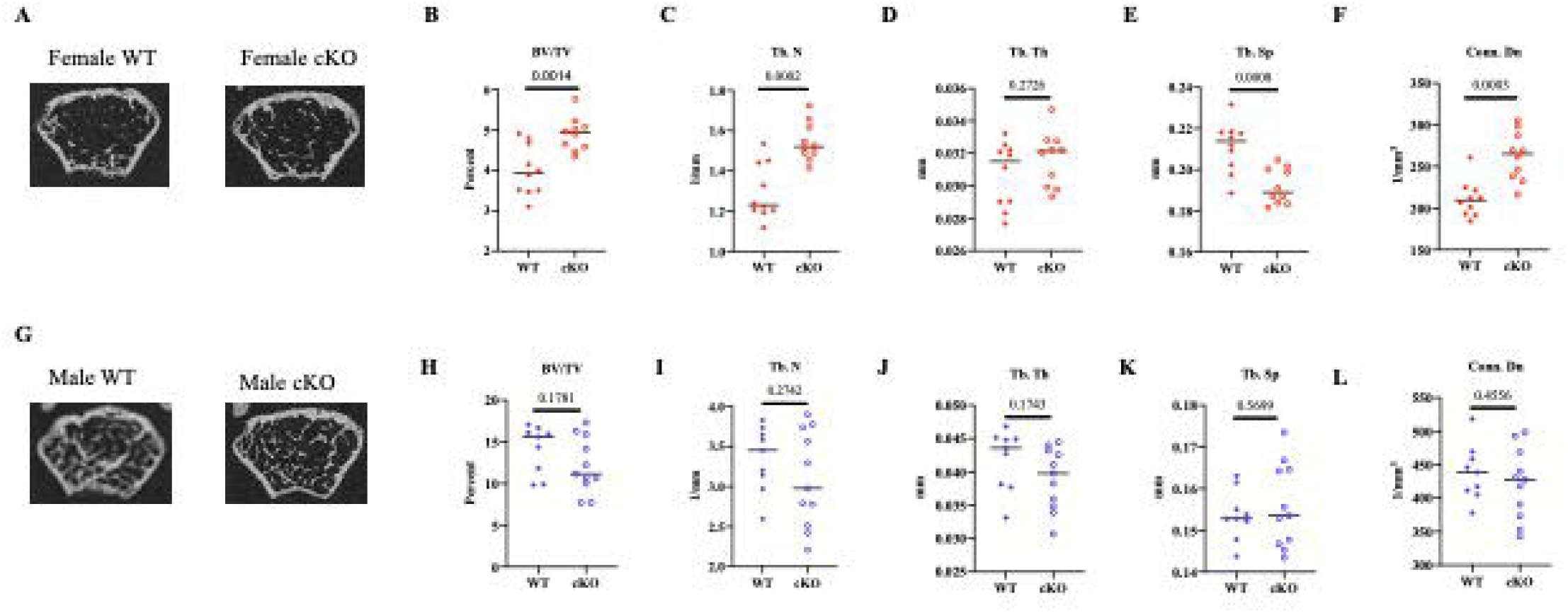
Female *Lsd1*cKO mice have increased bone mass at 3 months of age. Representative trabecular micro-CT images of female (A) and male (G) *Lsd1*WT and *Lsd1*cKO mice. measurements of (B and H) trabecular bone volume to total volume, (C and I) trabecular number, (D and J) trabecular thickness, (E and K) trabecular spacing, and (F and L) connective density are displayed. Female WT n=10, female cKO n=10, male WT n=9, male cKO n=11.

### Female Lsd1cKO mice have smaller osteoclasts

To determine if changes in *in vivo* osteoclast differentiation is contributing to increased bone mass in the Lsd1cKO mice, we performed histological analysis of paraffin embedded bones. Our analysis demonstrated that female Lsd1cKO mice do not have a difference in the number of TRAP positive osteoclasts but do have significantly smaller osteoclasts than female Lsd1WT mice (Oc.S/BS, 0.0002823±8.167e-005 decrease) (**Figure 2A-C**).

**Figure 2.**
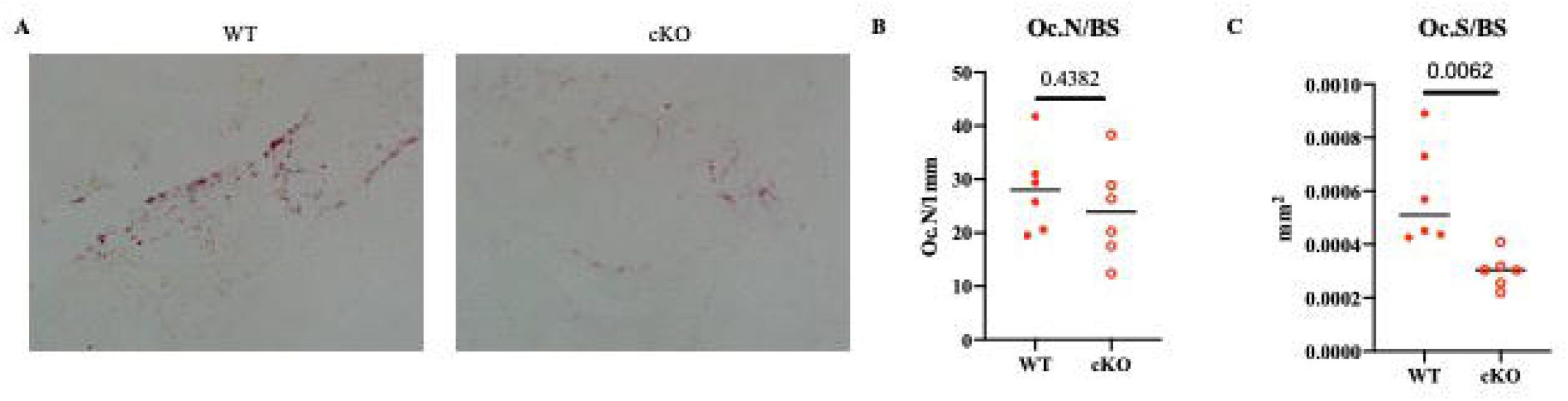
Female *Lsd1*cKO mice have smaller osteoclasts per bone surface. (A) Representative images of TRAP-stained bone slices from female femora of Lsd1WT and Lsd1cKO mice. Quantification of bone slices for (B) osteoclast number per bone surface and (C) osteoclast surface per bone slice. n=6 per genotype.

*In vivo* analysis of the male Lsd1cKO osteoclasts exhibit an interesting phenotype where there is no significant difference in the number of TRAP positive osteoclasts; however, the Lsd1cKO male mice have significantly larger osteoclasts (Oc.S/BS, 9.450e-005±3.928e-005) (**Figure S2A-C**). These data taken together with the uCT data suggests that LSD1 in myeloid cells does not regulate osteoclast differentiation in male mice. Therefore, in the rest of our analysis of LSD1 and osteoclasts, we focused on osteoclasts from female mice.

### Lsd1cKO osteoclasts have decreased differentiation

To determine the *in vitro* phenotype of osteoclasts derived from Lsd1cKO mice, we initially confirmed a significant decrease in LSD1 expression by qRT-PCR and western blot (**Figure 3A & B**). To measure osteoclast formation *in vitro*, we stimulated bone marrow monocytes with M-CSF and RANKL for 48 hours (day 2) and 72 hours (day 3). TRAP-stained images show that there are significantly less TRAP positive osteoclasts at day 2 and 3 of differentiation (**Figure 3C-E**). Additionally, at day 3 of differentiation, Lsd1cKO osteoclasts are significantly smaller in size compared to Lsd1WT (0.007700±0.0009610 smaller) (**Figure 3 F & G**).

**Figure 3.**
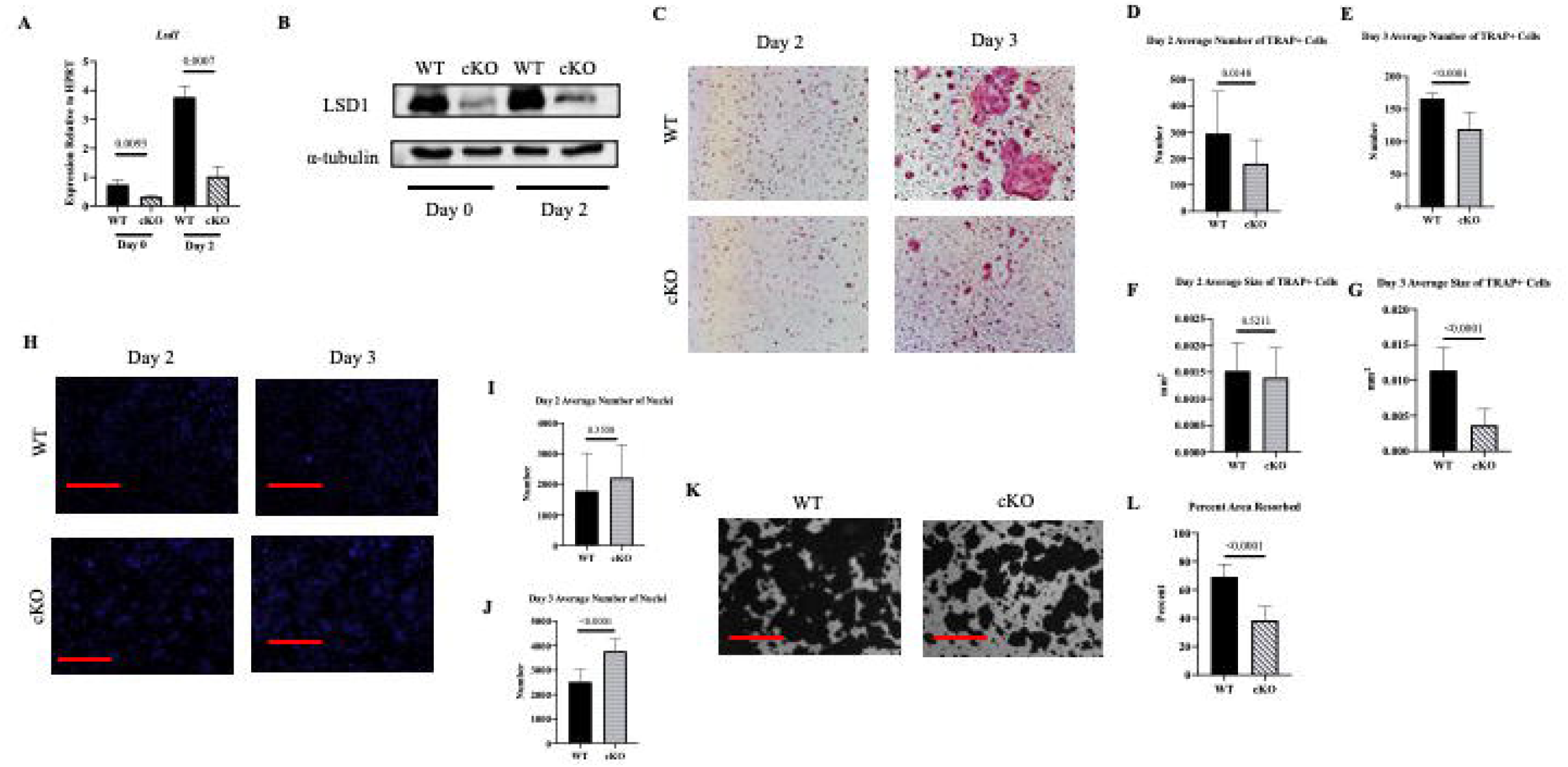
*Lsd1*cKO mice have less TRAP positive and smaller osteoclasts with decreased resorptive activity. Bone marrow monocytes from *Lsd1*WT and *Lsd1*cKO mice were stimulated with M-CSF and RANKL to induce osteoclast differentiation. (A) Representative TRAP-stained images of cells 48 hours (day 2) and 72 hours (day 3) after RANKL stimulation. (B) Average number of TRAP positive cells and (C) average size of TRAP positive cells at 48 hours. (D) Average number of TRAP positive cells and (E) average size of TRAP positive cells at 72 hours. (F) Representative DAPI-stained images of cells 48 hours and 72 hours after RANKL stimulation. Average number of nuclei present at 48 hours (G) and 72 hours (H). (I) Representative images displaying demineralization of calcium phosphate 120 hours (day 5) after RANKL stimulation. (J) Percent area of calcium phosphate resorbed.

To determine if the decrease in size of the TRAP positive osteoclasts was due to the inability to proliferate, osteoclast cultures were stained with DAPI to quantify the number of nuclei present. At day 2 of differentiation there was no significant difference between the number of nuclei and at day 3 of differentiation there were in fact more nuclei present in the Lsd1cKO cultures (**Figure 3 H-J**). This data suggests that osteoclast differentiation is not inhibited in the Lsd1cKO mice due to lack of precursor cells proliferating however, the decrease in size suggests they may be inhibited from fusing into multinuclear cells. To measure the ability of osteoclasts to demineralize we plated the osteoclasts on calcium phosphate coated plates to measure their ability to resorb mineral. As expected, based on their decreased size, Lsd1cKO osteoclasts were not able to effectively resorb as much mineral as their wildtype counterparts (30.64±2.892 decrease) (**Figure 3K & L)**. Lastly qRT-qPCR analysis shows that the loss of LSD1 results in a decreased trend in the osteoclast gene markers *Nfatc1, Dcstamp* and *Ctsk*, but no significant change in *c-Fos* expression (**Figure 6A-D**). These results signify that there is a decrease in genes that regulate osteoclast differentiation and activity in the absence of LSD1.

### Lsd1cKO osteoclasts have an increase in overall methylation at H3K4

To better understand the decrease in osteoclast differentiation in the Lsd1cKO mice, we determined the global methylation status of the two targets of LSD1: H3K4 and H3K9. Both day 0 (M-CSF only) and day 2 (M-CSF and RANKL) of osteoclast differentiation, there is an increase in mono-methylation of H3K4 in the Lsd1cKO osteoclasts, but no difference in mono-methylation of H3K9 at either day (**Figure 4E & F**). The decrease in methylation on H3K4 suggests that LSD1 inhibits genes to regulate osteoclast differentiation. Additionally, our qRT-PCR demonstrating that Lsd1cKO osteoclasts had no change in *c-Fos*, but a decrease in Nfatc1 expression suggests that LSD1 inhibits genes involved in osteoclast differentiation after the commitment phase.

**Figure 4.**
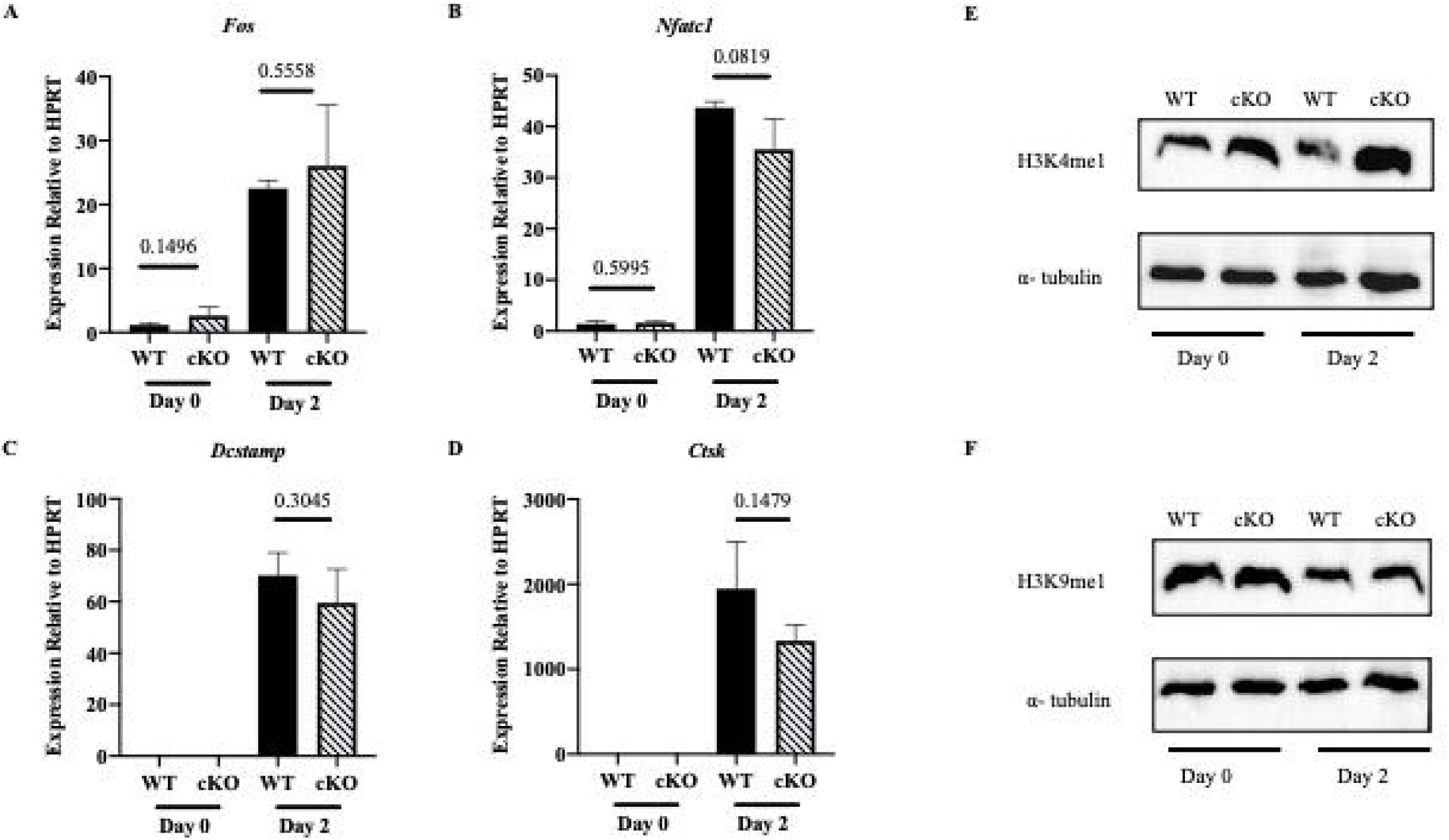
*Lsd1*cKO mice have decreased expression of osteoclast genes and increased mono-methylation at H3K4. (A-D) mRNA expression of osteoclast specific genes 0 hours and 48 hours after RANKL stimulation. Immunoblot analysis of mono-methylated H3K4 (E) and mono-methylated H3K9 (F) 0 hours and 48 hours after RANKL stimulation.

### LSD1 can form a complex with CoREST and HDACs 1 & 2

Many studies of LSD1 in other cell types. demonstrate that LSD1 forms a complex with the corepressor CoREST (RCOR1) and histone deacetylases HDAC1 and HDAC2. These interactions facilitate LSD1’s ability to successfully demethylase its target genes [15-17]. To determine if this was indeed the case in osteoclasts as well, we immunoprecipitated LSD1 in Lsd1WT osteoclasts and immunoblotted for CoREST, HDAC1 and HDAC2. We see that in osteoclast precursors (day 0), as well as mononuclear osteoclasts (day 2) LSD1 is present within an immune complex containing CoREST, HDAC1 and HDAC2. Additionally, when we attempt to immunoprecipitate LSD1 from Lsd1cKO osteoclasts, we see the loss of HDAC1 and HDAC2 binding (**Figure 5A**). These data demonstrate that LSD1 indeed forms this same complex in osteoclasts and CoREST along with the HDACs may be important for LSD1’s stability and demethylase function. To further investigate the importance of the HDACs in the complex, we sought to examine the acetylation status of H3K9, a target for deacetylation by class I HDACs [18, 19]. Interestingly, we see that with the loss of LSD1 expression there is an increase in overall acetylation of H3K9 (**Figure 5B**).

**Figure 5.**
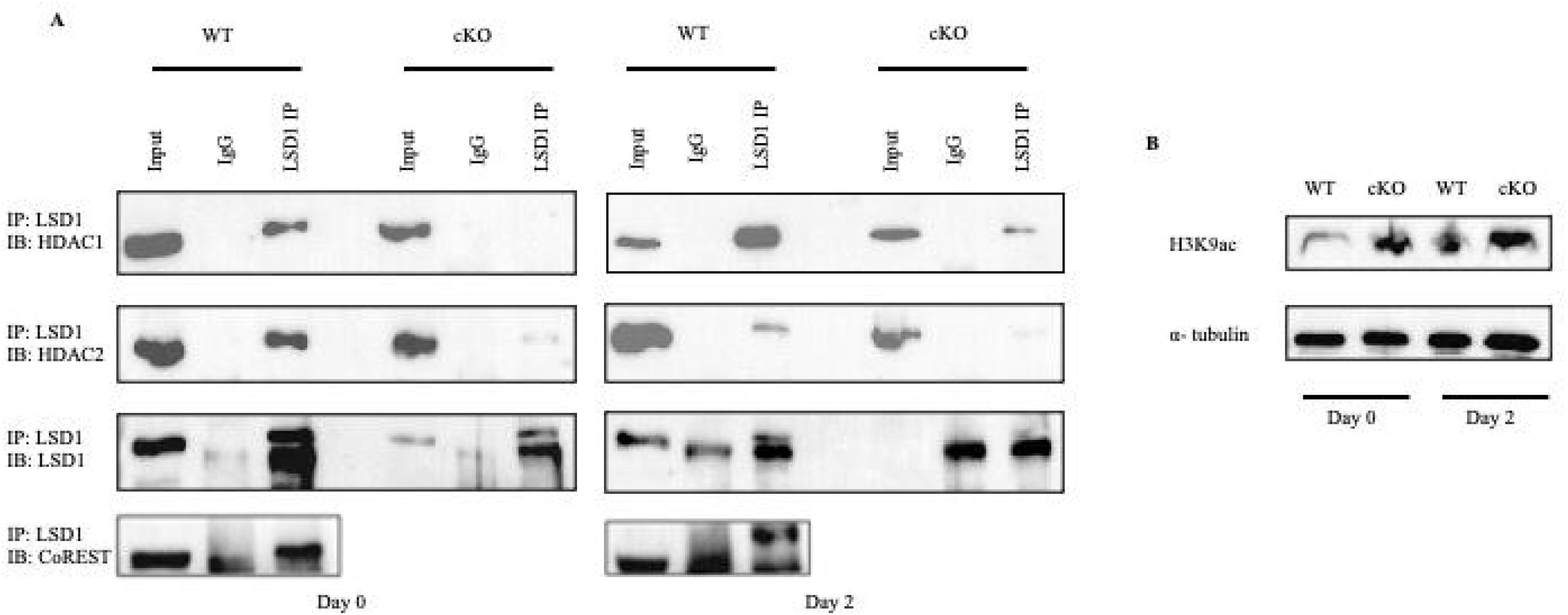
LSD1 can bind in a complex with HDAC1 and HDAC2. (A) LSD1 was immunoprecipitated and immunoblotting analysis was performed to look for the presence of HDAC1, HDAC2, LSD1 and CoREST at 0 hours and 48 hours after RANKL stimulation. (B) Immunoblot analysis of acetylated H3K9 0 hours and 48 hours after RANKL stimulation.

## Discussion

An imbalance in bone remodeling due to increased osteoclast differentiation ultimately leads to bone related diseases such as osteoporosis and periodontal disease [4]. A better understanding of the molecular mechanisms that regulate osteoclast gene expression will help us recognize how these cells promote bone loss and can uncover new possible drug targets [20]. Epigenetic factors have shown to be drug targets for regulating gene expression in a plethora of cell types [8]. One of these factors, specifically shown in cancers, is LSD1 [10-13]. However, to date, not much is known about the mechanisms by which LSD1 regulates gene expression during osteoclast differentiation.

In this study we show that conditional knockout of LSD1 in the myeloid lineage decreases the number and size of TRAP positive osteoclasts. To test these effects *in vivo*, we performed μCT to examine the effects on bone. We show that Lsd1cKO female mice have significantly more bone compared to their Lsd1WT littermates. Additionally, we show that Lsd1cKO female mice have smaller osteoclasts per bone surface. Interestingly, Lsd1cKO male mice have larger osteoclasts per bone surface compared to their Lsd1WT littermates; however, male Lsd1ckO had no significant skeletal phenotype compared to male Lsd1WT mice. Previous studies have shown that LSD1 interacts with the estrogen receptor α (ERα) in breast cancer cells and may suggest a mechanism for the fact that we only measured a significant skeletal phenotype in female Lsd1cKO mice [21]. We have attempted to confirm this in osteoclasts, however we have yet to find an antibody for ERα that recognizes the protein. Currently, we are confirming the LysM-cre phenotype by repeating our experiments in the monocyte population using Cfms-cre. If we find that there is a female only phenotype in the Cfms-cre as well, we hope to use an ovariectomized (OVX) mouse model to determine if female mice are more resistant to OVX induced bone loss in the absence of LSD1.

To determine the mechanism by which LSD1 may be regulating osteoclast differentiation, we investigated the loss of LSD1 and osteoclast gene expression. While not significant, the osteoclast gene markers *Nfatc1, Dcstamp* and *Ctsk* were downregulated in the LSD1cKO mice. Additionally, when looking at the LSD1 histone residue targets H3K4me1 and H3K9me1, we see an increase in the amount of H3K4me1 present in the Lsd1cKO osteoclasts. These data suggest that LSD1 may play a role in hindering osteoclast inhibitor gene expression via the removal of methylation from H3K4. Without LSD1, these genes are no longer repressed, and they can prevent osteoclast differentiation from occurring. However, further experiments such as chromatin immunoprecipitation need to be performed to confirm LSD1 target genes.

Much of our data agrees with a study determining the role of LSD1 in human osteoclasts [22]. This group showed that knockdown of LSD1 using shRNA in human osteoclasts suppresses differentiation and decreased pathological bone resorption in an arthritis mouse model [22]. They also demonstrated that RANKL stimulation induces LSD1 expression via the mTOR pathway [22]. While these results agreed with ours, the use of shRNAs in their study may cause off target effects and further studies would need to be performed to confirm their findings.

While much is still unknown about the role of LSD1 in osteoclast differentiation, it has been determined to be important in osteoblast differentiation. LSD1 regulates osteoblast differentiation by specifically targeting the genes *Wnt7b* and *Bmp2* [14]. The loss of *Lsd1* in both human mesenchymal stem cells (hMSCs) as well as mouse MSCs increased osteoblast differentiation and mineralization ability to increase bone volume [14]. Taken together, these data suggest that blocking LSD1 will decrease bone resorption and increase bone formation, making it an effective target for patients suffering from diseases such as periodontitis and osteoporosis. However, drug studies would need to be performed to confirm this.

To better understand how osteoclast differentiation may be inhibited, we are currently investigating the monocyte population in Lsd1WT and Lsd1cKO mice to determine if there are any differences in the cellular markers using flow cytometry. Though these experiments, we also hope to determine if the macrophage polarization ability of our monocyte populations differ between the two genotypes, which may suggest how LSD1 regulates monocyte, macrophage and osteoclast lineage commitment.

In conclusion, the loss of LSD1 in the myeloid lineage reduces the ability of osteoclasts to form and differentiate leading to an increase in bone volume in female mice.

## Figure Legends

**Supplementary Figure 1. Knockout of *Lsd1* does not affect cortical bone**. Representative cortical micro-CT images of female (A) and male (G) *Lsd1*WT and *Lsd1*cKO mice.

Measurements of (B and H) cortical bone volume to total volume, (C and I) cortical thickness, (D and J) mean total cross sectional bone area, (E and K) mean total cross sectional tissue area, and (F and L) mean total cross sectional tissue perimeter are displayed. Female WT n=10, female cKO n=10, male WT n=9, male cKO n=11.

**Supplementary Figure 2. Male *Lsd1*cKO mice have larger osteoclasts per bone surface**. (A) Representative images of TRAP-stained bone slices from male femora of Lsd1WT and Lsd1cKO mice. Quantification of bone slices for (B) osteoclast number per bone surface and (C) osteoclast surface per bone slice, n=6 per genotype.

