## Supplementary figures and images for "Female LSD1 Conditional Knockout Mice Have an Increased Bone Mass"

### Supplemental Figure 1

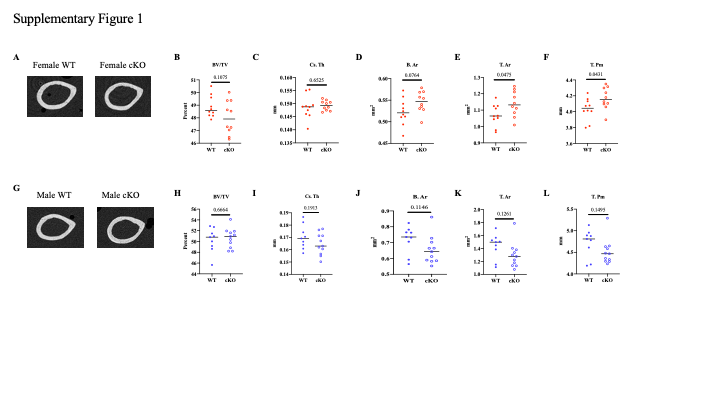

### Supplemental Figure 2

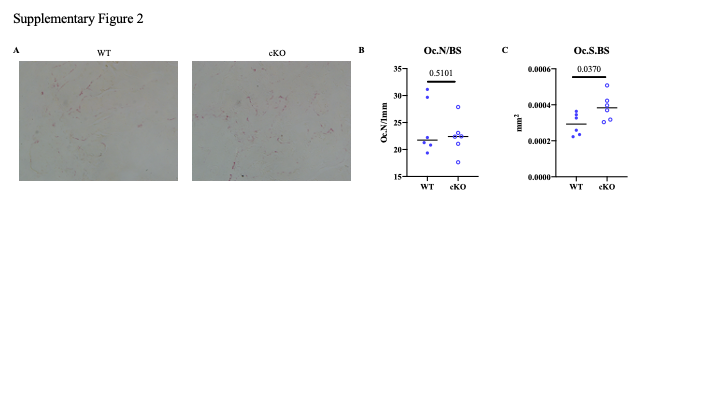
